# Reproducibility warning: The curious case of Polyethylene glycol 6000 and spheroid cell culture

**DOI:** 10.1101/793828

**Authors:** Simona Serrati, Chiara Martinelli, Antonio Palazzo, Rosa Maria Iacobazzi, Mara Perrone, Roberto Santoliquido, Quy K. Ong, Zhi Luo, Ahmet Bekdemir, Giulia Pinto, Ornella Cavalleri, Annalisa Cutrignelli, Valentino Laquintana, Nunzio Denora, Francesco Stellacci, Silke Krol

## Abstract

In this study we report about the reproducibility of three-dimensional cell culture of floating cell spheroids on PEG6000 treated cell culture dishes. Three-dimensional tumour spheroids or organoids present an interesting test platform for nanoparticulated drug delivery or nanoparticle toxicity. We tested the reproducibility of spheroid formation induced by the PEG coated surface. Interestingly we found that the results were different in a reproducible manner depending on the distributors of PEG6000.

Despite the nearly identical physicochemical properties of PEG6000 (MALDI-MS, NMR, FTIR, Triple SEC) with only minor differences, we observed only for one PEG6000 a highly reproducible formation of spheroids with different cell lines such as HT-29, HeLa, Caco2, and PANC-1. The surface coating with the different PEG6000 was studied by AFM. The surface coating as well as the physicochemical characterization showed only small differences in mass and hydrodynamic radius between the different PEGs. A direct coating of the cells with PEG from two distributors indicate that the spheroid formation in due to direct interaction of the polymer with the cell rather than by interaction of cells with the coated surface.

The experiments point out that for biological entities, such as cells or tissues, even very small differences such as impurities or batch-to-batch variations in the purchased product can have a very strong impact.

## Introduction

One of the major concerns about the quality of the research is the reproducibility either of the own data or also from other researchers [1] as Baker showed with the results from a survey answered by 1500 scientists. Numerous reasons have been identified for the failure to reproduce data, such as selective reporting or low statistical power or poor analysis. Especially in medicine, chemistry, and biology the reproducibility of data is an issue.

In the following, we present a study in which biology meets chemistry for medical applications where we identified another reason for failure to reproduce data. We performed a surface coating by adhesion with polyethylene glycol with a molecular weight of 6000 Da from different distributors. These surfaces should be cell repellent in order to induce cell growth in spheroids. These experiments were carried out in 2 different laboratories by 4 different operators repeatedly. We used a variety of cell lines. Three-dimensional (3D) tumor cell culture became an important tool as realistic test bed especially for nanoparticulated drug delivery, nanotoxicity as well as pharmaceutical drug testing as it mimics more closely the physiologic environment in a tumor in terms of accessibility, presence of extracellular matrix, and intercellular communication as compared to conventional monolayer (2D) cell culture [2]. In floating 3D spheroids, one can avoid the interaction and uptake of aggregated nanoparticles which can precipitate in long-term incubation on the cell surface in 2D cell culture conditions and induce artifacts in toxicity studies [3]. The impact on drug concentration and toxicity can be significant [4]. The presence of extracellular matrix and several layers of cells makes them a good model for the development and optimization of efficient intratumoral nanoparticulated drug delivery [5,6] and for drug penetration and diffusion studies [7–10]. Moreover, it has been demonstrated that large spheroids (>200 μm in diameter) form the three different regions of a tumor, such as a proliferating periphery region, a viable, but quiescent intermediate region, and a necrotic core [7,11,12]. The spheroids recapitulate *in vivo* tumor-like development patterns of avascular tumor nodules, in terms of morphology and growth kinetic properties [13–15].

Different techniques are available to grow small tumour spheroids. One of the simplest is the hanging drop cell culture for which a limited number of trypsinized cells from 2D cell culture are transferred with a limited amount of cell medium on the lid of the Petri dish. This lid is placed on a water-filled petri-dish. In the hanging drop, within 24 h, cells start clustering and form mostly small spheroids, one spheroid per drop [15].

In the following we will describe that by simply coating a culture dish with PEG6000 for 1 h at 37°C and cultivating the cells in these dishes. For PEG6000 from Carlo Erba we observed the formation of compact spheroids of varying size for different cell lines. The experiments were repeated with PEG6000 from other distributors (Merck, Sigma-Aldrich, Acros) and we observed that the other PEG6000 induced attached 2D growing cells. In order to identify the difference in the chemical composition, all PEGs were analysed in detail for physicochemical properties by NMR (nuclear magnetic resonance), FTIR (Fourier transform infrared spectroscopy), triple SEC (size exclusion chromatography), AFM (atomic force microscopy), and by MALDI-MS (matrix-assisted laser desorption/ionization mass spectroscopy). We found small differences in the molecular weight and viscosity. Finally, we determined that a direct cell-polymer interaction is responsible for the observed difference in cell growth by pre-incubating the cell with the polymer and then after washing deposit them on an uncovered culture dish. These experiments point out the importance of precise description of the purchased product in order to allow reproducibility.

## Materials and Methods

### Cells

Human colorectal cancer cell line HT29 was obtained from American Type Culture Collection. HT29 cells were cultured in McCoy’s 5a Medium Modified (Euroclone, Italy) with 10 % fetal bovine serum (FBS; Gibco, Thermo Fisher Scientific, Inc., Waltham, MA, USA), 1 % glutamine, and 1% penicillin/streptomycin. Cells were cultured in an incubator at 37 °C in an atmosphere containing 5% CO_2_. PANC-1 is a human pancreatic tumor cell line. HeLa is a cervical human adenocarcinoma cancer cell line obtained from American Type Culture Collection (ATCC). Hela cells were cultured in Eagle’s Minimum Essential Medium (EMEM) with 10% fetal bovine serum (FBS; Gibco, Thermo Fisher Scientific, Inc., Waltham, MA, USA) and 1 % penicillin and streptomycin. Caco-2 and HT29 are both human epithelial colorectal adenocarcinoma cell lines.

### Reagents

Polyethylene glycol; average molecular weight: 6000 Da (PEG6000) was purchased from Carlo Erba (C.E.), cat. n°: A192280010 (out of production); Sigma Aldrich (S.A.), cat. n°: 1546580; Merck Millipore, cat. n°: 8.07491; Carlo Erba as distributor for ACROS ORGANICS (COD. 192280010 LOT. A0398882). Polyethylene glycol 4000 (PEG4000) was purchased by Polichimica, Bologna, Italy, LOT no. 97001725.

### 3D cell culture

96-well plates were coated with a cell-repellent surface by exposing the wells for 1 hour at 37 °C with 200 μL of 3 % (w/v) of four different PEG6000, in Milli-Q water filtered with 0.22 μm filters. Then the PEG solution was removed and 12×10^4^ cells were seeded in a final volume of 150 μL for each well. Cells were cultured for 24 up to 96 hours at 37 °C, 5 % CO_2_. The experiments were reproduced by four different operators in two independent laboratories. For the experiments with PEG4000 (Polichimica), we performed the cell culture either after coating the 96-well plate with 3 % as with PEG6000 alone or with 5 % which is an equal molarity as the 3% PEG6000 solution. Next the plate was incubated with PEG4000 alone, or PEG4000 mixed with PEG6000 in a 1:1; 1:5 or 1:10 ratio. 1:5 is the ratio calculated from the spectrum in MALDI.

Additionally, HeLa and HT-29 cells were incubated for 5 mins in a 3 % PEG6000 solution from C.E. and S.A. For the experiment with short incubation, 12×10^4^ cells were deposited in an untreated 96-well plate and grown as described before for 48 h.

Cells were visualized by light microscopy using an OLYMPUS CKX41 microscope with a 10X/0,25 PHP objective.

### Nuclear Magnetic resonance (NMR)

^1^H NMR were performed with a Bruker AV-400 MHz after dissolving the PEG6000 in D_2_O. The NMR data were analyzed with MestreNova.

### Attenuated total reflectance-Fourier transform infrared (ATR-FTIR) spectroscopy

The ATR-FTIR spectra were recorded using the IR spectrometer Thermo Fischer Scientific Nicolet 6700. 2 mg of the PEG samples were placed on the diamond crystal. The ATR measurement mode was used covering the range of 4000-600 cm^−1^. 64 scans were accumulated with a resolution of 4 cm^−1^ for each measurement. The spectra were processed using OMNIC spectra software to subtract the background and adjust baseline.

### Matrix assisted laser desorption ionization-time of flight mass spectrometry (MALDI-TOF-MS)

To prepare the sample for MALDI-TOF, the PEG samples were dissolved in methanol to make a 5 mg/mL solution. α-Cyano-4-hydroxycinnamic acid (CHCA) was used as the matrix and was dissolved in methanol with a concentration of 10 mg/mL. 10 μL of PEG solution was then mixed well with 10 μL matrix solution in an Eppendorf tube. 2 μL of the mixture solution was spotted onto the Bruker stainless steel target and completely dried under vacuum. MALDI-TOF mass spectra were obtained using a Bruker AutoFlex spectrometer. The mass spectra were measured with positive ionization and linear mode with laser power at 25 % attenuation. The resulted spectra were processed and plotted using FlexAnalysis software.

### Electrospray ionization time-of-flight mass spectroscopy (ESI-TOF MS)

#### Triple size exclusion chromatography (SEC)

The size exclusion chromatography was performed using the OMNISEC Triple Detection system produced by Malvern Panalytical, equipped with refractive index (RI), viscosimeter, low angle light scattering (LALS) and right-angle light scattering (RALS). The eluent was 0.1 M NaNO_3_ solution, flow rate 0.6 mL/min, column set G2500PWXL + G3000PWXL and the dn/dc applied was 0.14. The samples were dissolved in a small volume of the eluent and 100 μL of this solution was injected.

#### Atomic force microscopy (AFM)

Samples for AFM analysis were prepared by dissolving PEG in Milli-Q water to a concentration of 3 % (w/vol). A drop of PEG solution was placed on a polystyrene Petri dish (Iwaki, 1000-035, non-treated) for 1 h at 37 °C. Then the solution was removed, and Milli-Q water was added. Hydrated samples were then mounted into the AFM liquid cell. AFM measurements were carried out using a Multimode/Nanoscope V system (Bruker). AFM images were acquired in contact mode in liquid (MilliQ water) using commercial Si3N4 cantilevers (DNP-10 Bruker, k=0.24 N/m). For some samples PEG solution was not removed and AFM measurements were performed in PEG solution. No significant differences were observed between images acquired in Milli-Q water or PEG solution. Data were analyzed with Gwyddion software.

## Results

### Spheroid culture

We prepared the surface coating of the 96-well dish with PEG6000 from different distributors using always the same concentration (3 %) and incubation conditions (37 °C, 1 h). As shown in figure 1, only the PEG6000 from C.E. provided a cell-repellent surface necessary to form tight and well-defined floating 3D spheroids (fig.1A) in the solution rather than a 2D cell layer.

**Figure 1.**
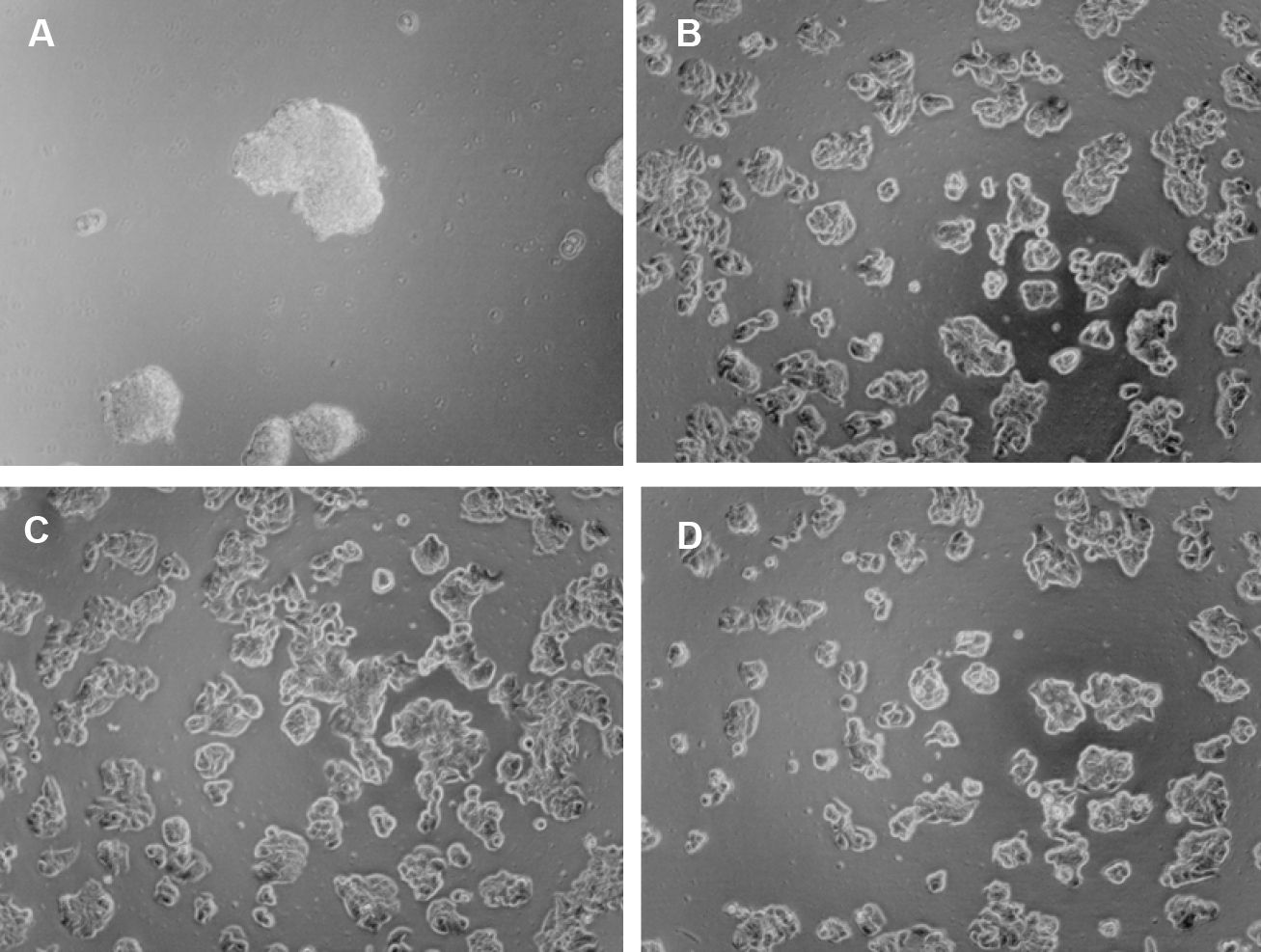
Microscopic images of HT29 cells on surfaces (1 h; 37 °C) treated with PEG6000 from A) C.E.; B) Acros (C.E.); C) Merck; and D) S.A.. The images were recorded 48 h after cell plating.

In order to exclude that it was an exceptional result due to properties of the cell line HT29, we performed the same experiments with different cell lines such as Caco-2, PANC-1, or HeLa. As depicted in figure 2, all tested cell lines showed the same spheroid formation when PEG6000 from C.E. was used to coat the well plates. On the PEG6000 products from other distributors we observed cell attachment and 2D culture as observed for HT29 (data not shown).

**Figure 2:**
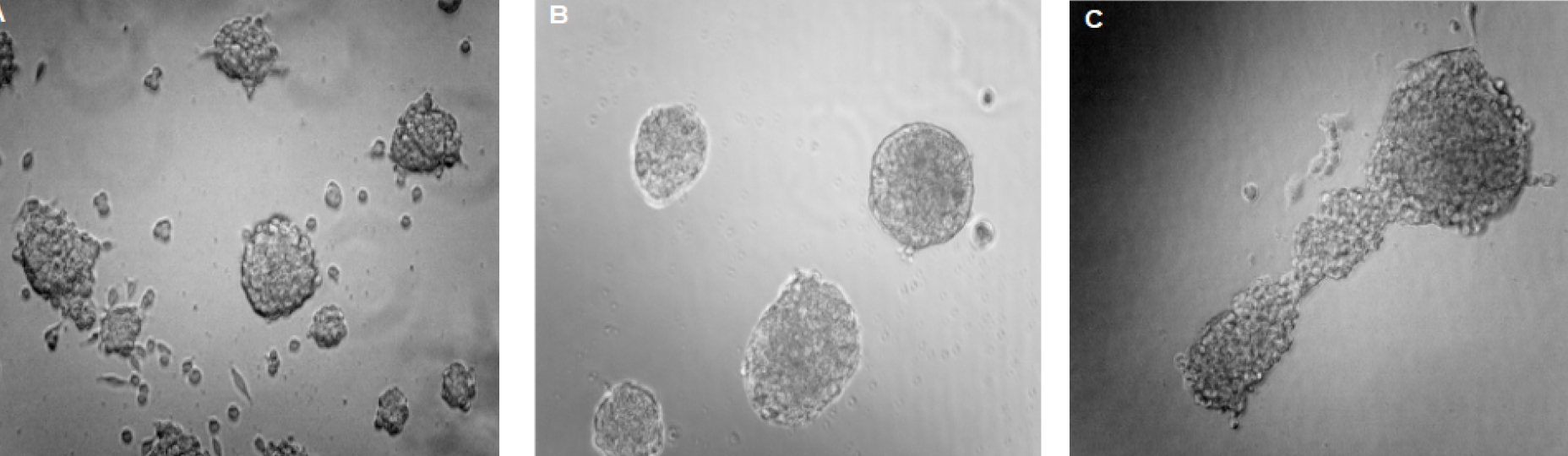
Micrographs of (A) HeLa, (B) PANC-1; and (C) Caco-2 cells imaged 72 h after seeding on PEG60000 (C.E.) pre-treated surfaces.

The difference in cell behaviour on surfaces pre-treated with PEG6000 from different distributors led us to determine in-depth the physicochemical properties of the purchased PEG.

### Physicochemical characterization of the PEG6000

First, we measured the mass of the different PEGs either by MALDI-MS, triple SEC, or ESI-TOF-MS. MALDI spectra of PEG6000 from different distributors showed a small difference (fig. 3). While average mass for PEG6000 from C.E. and Merck is around 6000 Da, as expected, the average mass for PEG from S.A. and Polichimica is closer to 7000 Da. The only difference we detected in the mass spectrum of PEG from C.E was an additional peak at 4000 Da with a relative ratio of 1:5 to PEG6000 (calculated by peak height).

**Figure 3.**
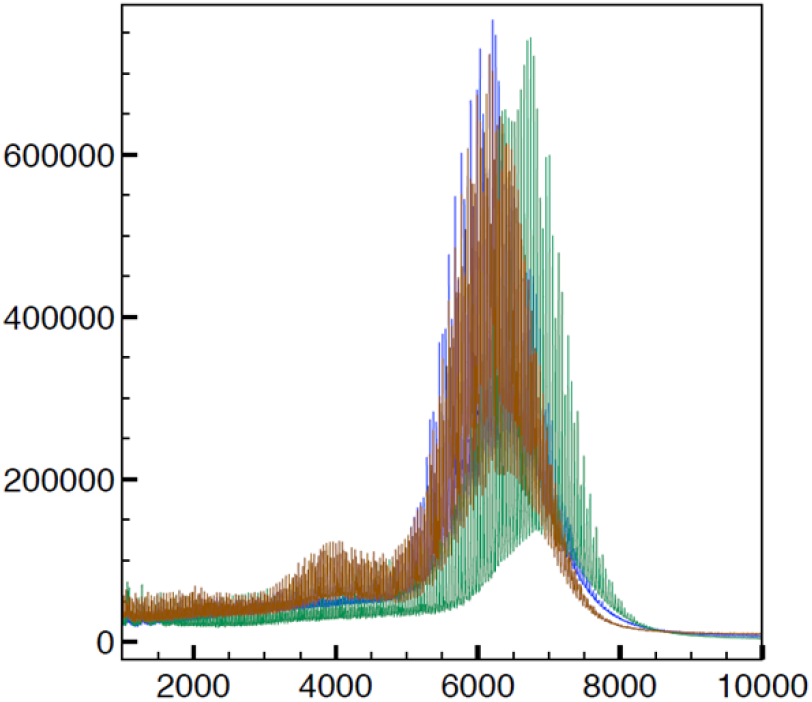
(A) MALDI measurements to determine the mass of PEG 6000 produced by S.A. (green), Merck (blue), C.E. (red).

We performed FTIR and NMR measurement to determine the nature of the lower molecular weight peak. As it can be seen in figure S1 in Supporting Information (SI) the spectra in FTIR are substantially similar for the 3 tested PEG solutions. The measurements in ^1^H-NMR (fig. S2 in SI) confirmed that the chemical identity of the molecules is the same. From the NMR and FTIR measurements it was clear that the second mass peak is PEG4000 in the C.E. PEG. In order to understand if this lower weight PEG is responsible for the cell-repellent properties we coated the surface with PEG4000 (Polichimica) alone using different concentrations (3 %, 5 %), or in combination with different ratios of PEG6000 (from S.A. or Merck): PEG4000 (1:1; 1:5; 1:10) and using different mixing procedures (fig.4)..

**Figure 4.**
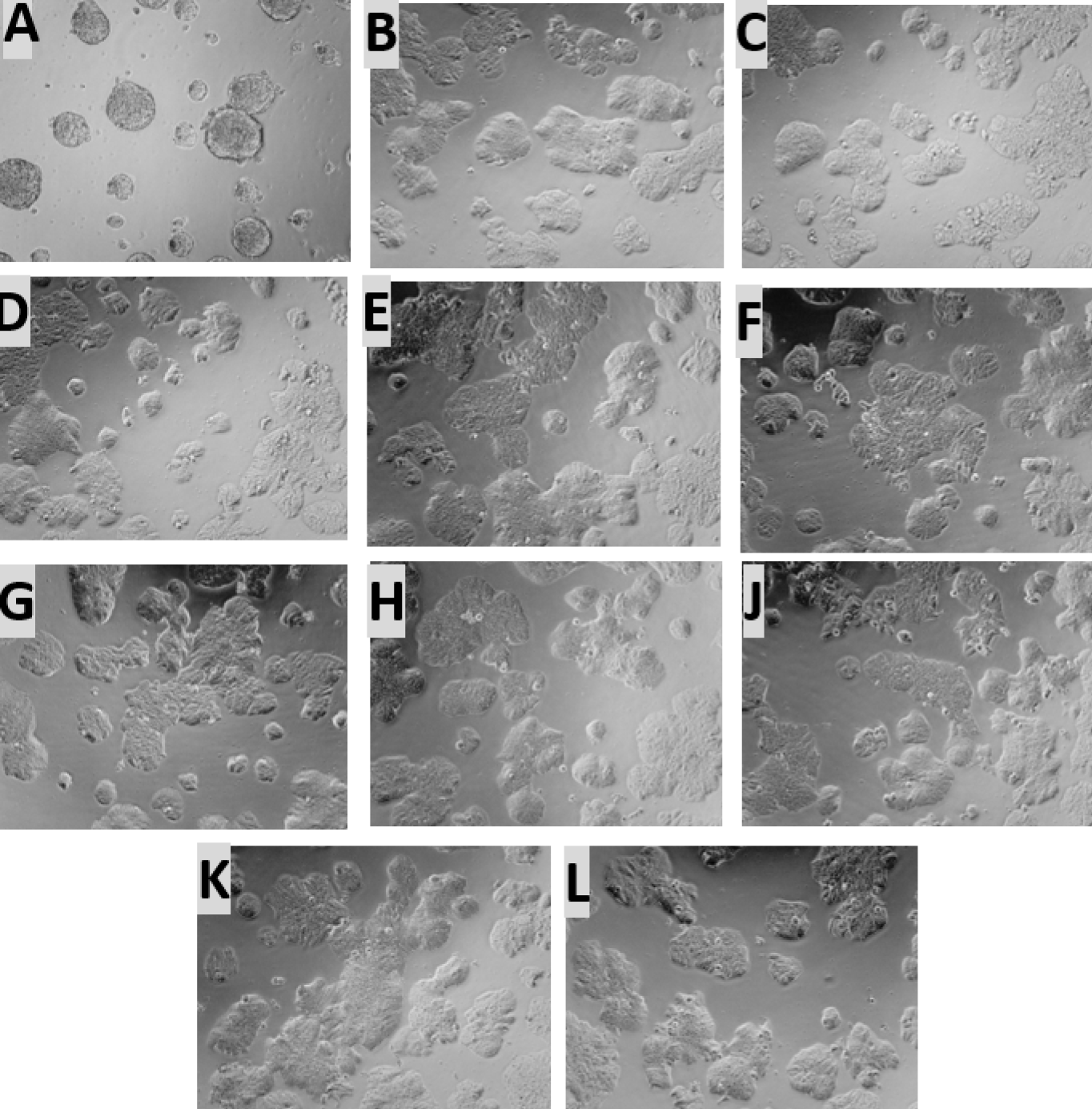
HT29 imaged by transmission light microscopy 96 h after plating in a 96 well plate treated for 1 h at 37 °C with (A) PEG6000 from C.E., (B) PEG4000 (4000), (C) PEG6000 from S.A. (D) a mixture of 4000/S.A. 1/5 PRE diluted, (E) a mixture of 4000/S.A. 1/5, (F) mixture of 4000/S.A. 1/10, (G) PEG6000 from MERCK, (H) 4000/MERCK 1/5 PRE diluted, (J) 4000/MERCK 1/5, (K) 4000/MERCK 1/10, and (L) as control HT29 cells on an untreated surface.

As it can be seen in the micrographs in figure 4, if PEG4000 (Polichimica) is added in different ratios, as it was measured for the mass spectrum of PEG from C.E. to the PEG6000 from other distributors the cells still grow in 2D and not in spheroids.

Cells are known to be responsive to micro- and nano-structured surfaces (e.g. [16]). In order to understand if a stable secondary structure of the polymer such as a random coil that are deposited on the surface in the coating process is responsible for the 3D spheroids, we performed triple size exclusion chromatography (SEC) measurements. These SEC measurements combining measurements of refractive index, viscosity, and right angle light scattering (RALS) allow to determine the molecular weight and the viscosity, and get information on the structure of the polymers. The results are summarized in Table 1.

**Table 1.** Triple SEC measurement (refractive index, viscosity; RALS) with PEG6000 from S.A. (Sigma), C.E. (Carlo Erba), and Merck.

. Molecular weight (Mw), the intrinsic viscosity ( $\eta$ ) and the hydrodynamic radius (Rh)
| Sample | Inj | Mn (Da) | Mw (Da) | Mw/Mn | $[\eta]$ (dl/g) | Rh (nm) | Recovery % |
| --- | --- | --- | --- | --- | --- | --- | --- |
| A6863 Sigma | 1 | 6275 | 6313 | 1,006 | 0,17 | 2,56 | 94,74 |
|  | 2 | 6318 | 6371 | 1,009 | 0,169 | 2,57 | 94,54 |
|  | average | 6296,5 | 6342 | 1,0075 | 0,1695 | 2,565 | 94,64 |
| A6864 Carlo Erba | 1 | 5821 | 5977 | 1,027 | 0,164 | 2,48 | 95,38 |
|  | 2 | 5864 | 5931 | 1,011 | 0,161 | 2,47 | 95,32 |
|  | average | 5842,5 | 5954 | 1,019 | 0,1625 | 2,475 | 95,35 |
| A6865 MERCK | 1 | 6020 | 6071 | 1,008 | 0,166 | 2,51 | 95,71 |
|  | 2 | 6017 | 6105 | 1,015 | 0,166 | 2,51 | 95,53 |
|  | average | 6018,5 | 6088 | 1,0115 | 0,166 | 2,51 | 95,62 |

As it can be seen in table 1, SEC confirms the slightly lower molecular weight of PEG6000 from C.E. and Merck, as already measured by MALDI. Moreover, PEG6000 from C.E. had also the lowest intrinsic viscosity and a smaller hydrodynamic radius which could be indicative for a more compact, coiled structure.

Next, we investigated if the small differences in hydrodynamic radius measured in triple SEC have an influence on the surface coating of PEG on the cell culture dish by visualizing the surface in atomic force microscopy (AFM). The dish was incubated with 3 % PEG solution. Then the solution was replaced by water and the surface was imaged by atomic force microscopy (AFM) in contact mode.

In the AFM micrographs (fig. 5) we observed no remarkable differences in the surface pattern, island size or height or distance between the polymer islands for surfaces coated with PEG (C.E.) with respect to the others. As also the surface was not responsible for the spheroidal growth of the cells and because there was no washing between incubation with PEG solution and addition of the cell suspension we tested if residual PEG can be the reason for the cell growth. So we incubated the cells briefly in 3 % PEG6000 from C.E and from S:A. and seeded the cells on an uncoated 96-well plate. In figure 8 the cells were imaged 48 h after seeding.

**Figure 5:**
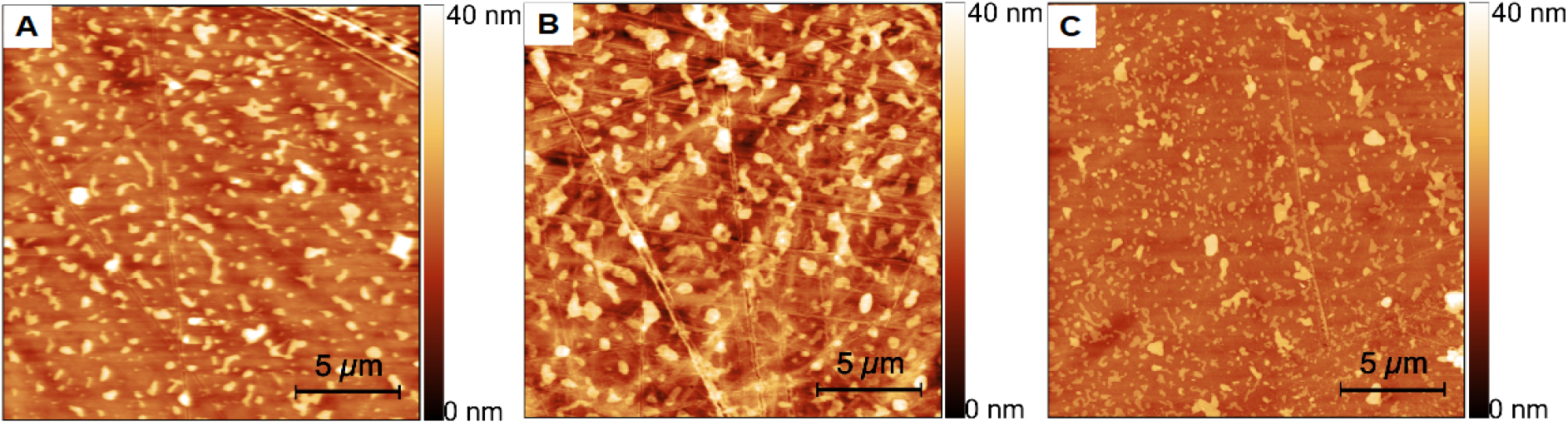
AFM micrograph of Petri dish surfaces incubated for 1 h with 3 % PEG6000 from (A) C.E.; (B) Merck; and (C) S.A. The images were recorded after replacing the PEG solution by Milli-Q water.

The micrographs show that both cell lines, HT-29 and HeLa gave round single cells if incubate in 3 % PEG6000 (C.E.) for 5 mins before seeding in the cell culture plate. while no differences between untreated control cells and PEG6000 (S.A.) incubated cells could be seen. This indicates that the reason for the spheroidal growth in PEG6000 from C.E. is due to a direct cell-polymer interaction rather than an effect of the surface coating as can be seen in the figure 6.

**Figure 6:**
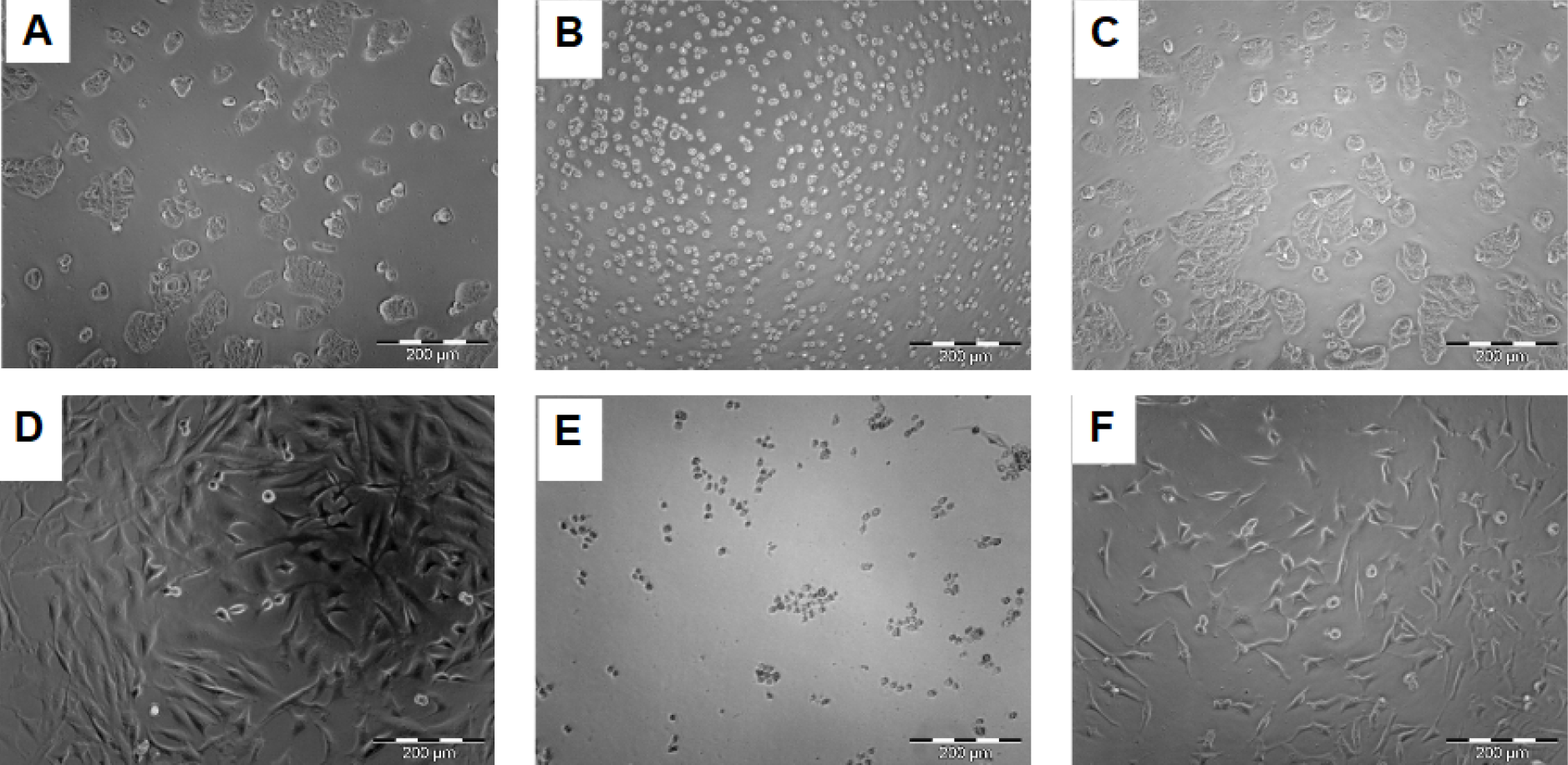
Micrographs of HT-29 (A-C) and Hela (D-F) cells without treatment (A, D), or incubated for 5 mins in PEG6000 from C.E. (B, E) and S.A. (C, F).

## Conclusion

In the present study, we tested the cell repellent properties of PEG6000 from different distributors. While we found no significant differences in the chemical profile of the different materials, except that for two PEG6000 the molecular weight was closer to 7000 Da and in one case there was an impurity of 4000 Da PEG, we observed a significant difference in the biological response in terms of cell growth. This highlights the significance for a precise description of the used material in order to allow reproducibility.

## Acknowledgement

The author(s) received no specific funding for this work.

